# Elaborate pupils in skates may help camouflage the eye

**DOI:** 10.1101/483172

**Authors:** Sean Youn, Corey Okinaka, Lydia M Mäthger

**Affiliations:** Marine Biological Laboratory, Bell Center, Woods Hole, MA, USA; Wiess School of Natural Sciences, Rice University, Houston, TX, USA; Biological Sciences Division, University of Chicago, Chicago, IL, USA

**Keywords:** crypsis, spatial frequency, background, vision

## Abstract

The little skate *Leucoraja erinacea* has elaborately shaped pupils, whose characteristics and functions have not been studied extensively. It has been suggested that such pupil shapes may camouflage the eye; yet, no experimental evidence has been presented to support this claim. Skates are bottom-dwellers that often bury into the substrate with their eyes protruding. If these pupils serve any camouflage function, we expect there to be a pupillary response related to the spatial frequency (“graininess”) of the background against which the eye is viewed. Here, we tested whether skate pupils dilate or constrict in response to background spatial frequency. We placed skates on background substrates with different spatial frequencies and recorded pupillary responses at three light intensities. In experiment 1, the skates’ pupillary response to three artificial checkerboards of different spatial frequencies was recorded. Skates responded to changing light intensity with pupil dilation/constriction; yet, their pupils did not change in response to spatial frequency. In experiment 2, in which skates could bury into three natural substrates with different spatial frequencies, such that their eyes protruded above the substrate, the pupils showed a subtle but statistically significant response to changes in substrate spatial frequency. Given the same light intensity, the smaller the spatial frequency of the natural substrate, the more constricted the pupil. While light intensity is the primary factor determining pupil dilation, these experiments are the first to show that pupils also change in response to background spatial frequency, which suggests that the pupil may aid in camouflaging the eye.

## Introduction

The study of camouflage has received an enormous amount of attention in the past two decades, with research encompassing a wide array of study organisms, from animals that actively select habitats to accommodate their fixed body colors and patterns, to those who have changeable camouflage and can adjust their colors and patterns to effectively blend in with their surroundings (e.g., Cott, 1940; Merilaita, 1998; Cuthill et al., 2005; Stevens et al., 2006; Mäthger et al., 2009; Stuart-Fox and Moussalli, 2009; Zylinski and Johnsen, 2011; Hanlon and Messenger, 2018). Interestingly, the majority of this research has concentrated on the animal’s body camouflage; what has received scant attention is the question of whether and how animal eyes may be camouflaged, and what role the pupils may play.

Pupils restrict the optical aperture in the eyes of vertebrates and invertebrates. Most pupils respond to light by constricting. This carefully regulates the amount of light entering the eye and it is a crucial component in optimizing resolution and sensitivity in the retina (Wilcox and Barlow, 1975; Hammond and Mouat, 1985; Land and Nilsson, 2012; Douglas 2018). A round, dilated pupil is often a very conspicuous structure that stands out, possibly giving away the location of a lurking predator, or camouflaging prey animal. Circular pupil shapes can be very conspicuous (e.g., Thresher, 1977) because they present clear edges, and it is well known that contrasting edges may be visually easy to detect (Troscianko et al., 2009). Some animals have unusually shaped pupils, such as the mobile W-shaped pupil of cuttlefish, or the U-shaped pupils that are seen in some skates, rays, fish and squid. Several publications state that these pupils are found in species that spend significant amounts of time camouflaging on the substrate, and that these pupils might help camouflage the eye (Muntz, 1977; Douglas 2018; Douglas et al., 1998; 2002; 2005). By giving the pupil outline a less regular shape, allowing the eye to better blend into its surroundings, the likelihood of visual edge detectors recognizing the shape as belonging to an eye would be reduced. However, this claim has never been put to the test.

There are only a handful of publications that report specific eye adaptations and pigment markings that reduce the conspicuousness of the eye (Barlow, 1972; Nilsson and Nilsson, 1983; Neudecker, 1989). Yet, for some animals, eye contact is actively sought, as it is an important indicator of emotional state and intention (Burger et al., 1992). Laboratory and field studies have shown that gaze direction (i.e., whether eye contact is made or not) is an important feature perceived by animals and that it determines the observer’s behavior (Gagliardi et al., 1976; Hennig, 1977; Burger and Gochfeld, 1981; Burghardt and Greene, 1988). In a predator-prey context, eye contact may mean detection is more likely. Therefore, when trying to locate a camouflaged predator or prey, identifying the eyes may be of utmost importance; and, from the perspective of the animal that hides from potential predators or prey, hiding the eyes should be paramount. This should be even more important for animals that generally bury in the substrate with only their eyes protruding, as is true of the little skate *Leucoraja erinacea*.

*Leucoraja erinacea* has a very unusual pupil shape that, when seen under bright light, consists of a series of frills (Fig. 1). Under low light, the pupil opens to form an almost circular shape. To date, there are no descriptions in the literature regarding the function of this pupil. *L. erinacea* is a near-shore/shallow-water species that inhabits depths to about 90 m at maximum (Robins and Ray, 1986). Their eyes lack cone photoreceptors and therefore color vision, and their rods have a maximum spectral absorbance at around 500 nm (Dowling and Ripps, 1970; Cornwall et al., 1989). Batoid elasmobranchs implement a range of feeding strategies, including continuous feeding or foraging, ambush predation and filter feeding. Skates are important predators in benthic and demersal communities, preying mostly on fish and invertebrates, and, depending on life history stage, they adopt mixed feeding strategies, including ambushing teleost fish and squid, and searching for smaller invertebrates (McEachran and Musick, 1975; Ajayi, 1982; Ebert *et al.*, 1991; Orlov, 1998, Mabragaña & Giberto 2007, Jacobsen & Bennett 2013).

**Fig. 1.**
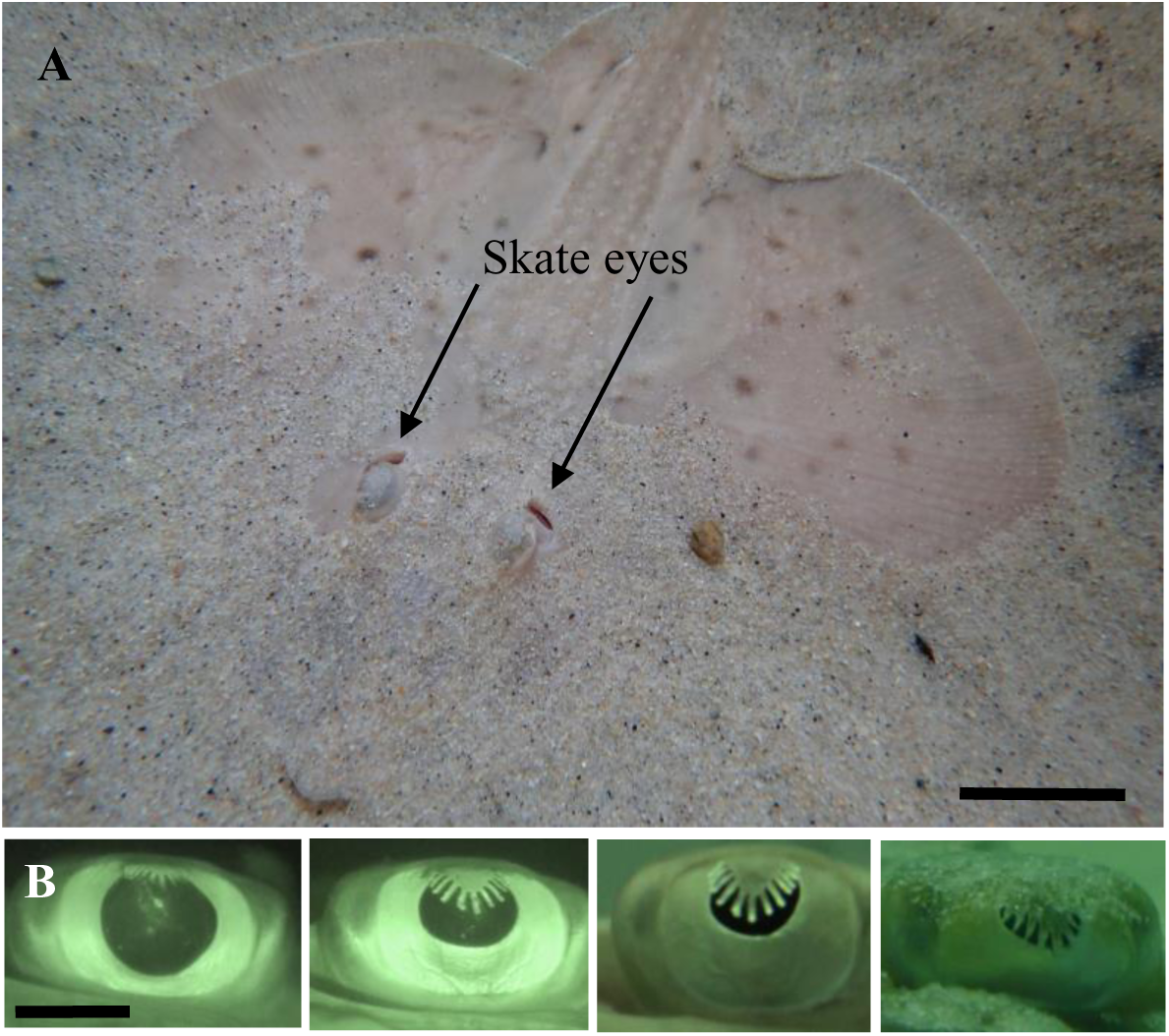
(A) Skate *Leucoraja erinacea* partially buried in sand. Scale bar: 8 cm. (B) close-ups of skate eyes showing range of dilation/constriction states from fully dilated (left) to near-constricted, revealing elaborate, finger-like shape. Scale bar: 1 cm.

Importantly, while light intensity is the most obvious driving force of the pupillary reflex, it is known that light is not the only factor influencing pupil dilation and constriction (Hess and Polt, 1964; Muntz, 1977; Messenger, 1981; Gamlin et al., 1998; Barbur et al., 2002; Einhäuser et al., 2007; Bradley et al., 2008; Douglas, 2018). For this reason, it may not be surprising to find that animals, especially those with elaborate pupil shapes, actively using their pupils for non-vision related tasks, such as camouflage.

The goal of our study was to test whether the pupil may function in camouflaging the eye. We conducted two experiments: In the first, we presented skates with checkerboards of different spatial frequencies. This technique has been extensively used to study cuttlefish and fish camouflage (e.g., Ramachandran et al., 1996; Mäthger et al., 2006; Kelman et al., 2007; Barbosa et al., 2008; Allen et al., 2010; Buresch et al., 2015). By presenting animals with backgrounds that have known spatial frequencies, such as a black and white checkerboard, an animal that has an innate camouflage response will do its best to attempt to respond to this checkerboard accordingly, and this type of behavioral assay has proven to be very robust. In the second experiment, we presented skates with natural substrates that allowed animals to bury with only their eyes protruding. If the pupil in *L. erinacea* fulfills a camouflage function, we would expect to see the pupil change in response to changes in spatial frequency of the surrounding habitat.

## Materials and Methods

### Animals

The little skate *Leucoraja erinacea* is a local animal off the coast of Woods Hole, MA. Animals are regularly collected, as well as raised in captivity. All animals used for these experiments were adults, measuring between 25-43 cm (nose to tip of tail). They were kept at 12-15ºC. For the first experiment, 8 animals (2 females, 6 males) were kept in separate partitions, measuring length: 75 cm x width: 43 cm x height: 40 cm. Since these experiments took several weeks to complete, and skates are better kept in larger space, we modified our holding tanks for the second experiment. For the second experiment, 14 skates (6 females, 8 males) were kept as a group in a large holding tank (1.2 m x 3.6 m; 40 cm height). Individual skates were recognized by sex (males and females are easy to tell apart), specific body markings, and size. Skates were fed five times a week (variation of squid, butterfish and capelin), and all were cared for and experiments were conducted in accordance with the regulations of the MBL Institutional Animal Care and Use Committees.

### Experimental setup

The experimental setup consisted of a glass tank that was placed inside an improvised dark room constructed from an ice fishing tent (Shappell DX4000) surrounded by metal walls and black plastic sheeting that ensured there was no stray light entering the experimental setup. All video recording was done using a Sony HDR-XR550VEB video camera, equipped with night-shot, so that skate eyes could be recorded under low light. All experiments were taken between 9am and 5pm. Experiments 1 and 2 required different materials, and methods were therefore slightly different.

*Experiment 1: Checkerboard experiment*. The experimental tank measured 90 cm length, 30 cm width, 40 cm height, lined on the outside (all four tank sides and a lid, which was placed on top of the tank during experiments) with a white diffuser filter (Lee Filters Half White Diffusion) and a blue-green filter (Roscolux #374), to give skates a diffuse light field similar to a natural underwater habitat, and, additionally, shield animals from unwanted visual stimuli (e.g., experimenter moving). The light source, placed approximately 15 cm above the tank, consisted of two side by side custom-made light boxes that illuminated the entire tank evenly (each box was equipped with a Phillips 60W compact fluorescent bulbs and a Lee Full White Diffusion filter). The light intensities were measured with an Extech EasyView 30 light meter with an irradiance probe, which was placed on the bottom of the tank to collect downwelling light. The light intensity across the tank varied by less than 5%; at full illumination, the intensity was approximately 700 l× (note, using a lux meter, instead of a full spectrum radiometer, is convenient and sufficient for these experiments, since skates are monochromats with a maximum sensitivity at around 500 nm, which overlaps with the maximum sensitivity of the Easytech light meter). Three light intensities were created using neutral density filters (combination of 0.15ND and 0.6ND, Lee Filters). In a separate experiment, we verified the light intensities needed to obtain a dilated, semi-constricted, and near-constricted pupil (note, the light-emitting pupil area of a fully constricted pupil is difficult to analyze, therefore we chose a pupil constriction state that was near-constricted). These were 0.68 l×, 10.9 l× and 43.75 l×, respectively. The pupil dilation and constriction states, and required light intensities were obtained during an unrelated experiment.

We created three black and white (i.e., 0,0,0 and 255,255,255 RGB) checkerboard substrates of different spatial frequencies (Fig. 2A). To create the three spatial frequencies, we measured the anterior-posterior dimension of all skates’ eyes. These dimension (distance) values were used to make the checkerboards. The large checkerboard had a check size of 100% of this distance; the medium checkerboard had a check size of half; the small checkerboard had a check size of 25% of this distance. Not all skates had the same eye size; thus we made three sets of checkerboards to accommodate all skate eye sizes. The skates were organized into three groups based on eye size. Group 1: four skates had the same eye size to within less than 1 mm (average 6.96 mm; 0.24 s.d.). Their three checkerboards had check sizes of 7 mm, 3.5 mm and 1.75 mm, respectively. Group 2: Three skates had larger eyes (average 7.87 mm; 0.16 s.d.), and their three checkerboards had a check sizes of 8 mm, 4 mm and 2mm, respectively. Group 3: One animal had the largest eyes, and its three checkerboards had check sizes of 10 mm, 5 mm and 2.5 mm, respectively. All checkerboards were printed on regular matte printing paper and laminated to be waterproof (matte laminate). Since all checks were made up of the same black and white shades, overall reflectance of all substrates was the same.

**Fig. 2.**
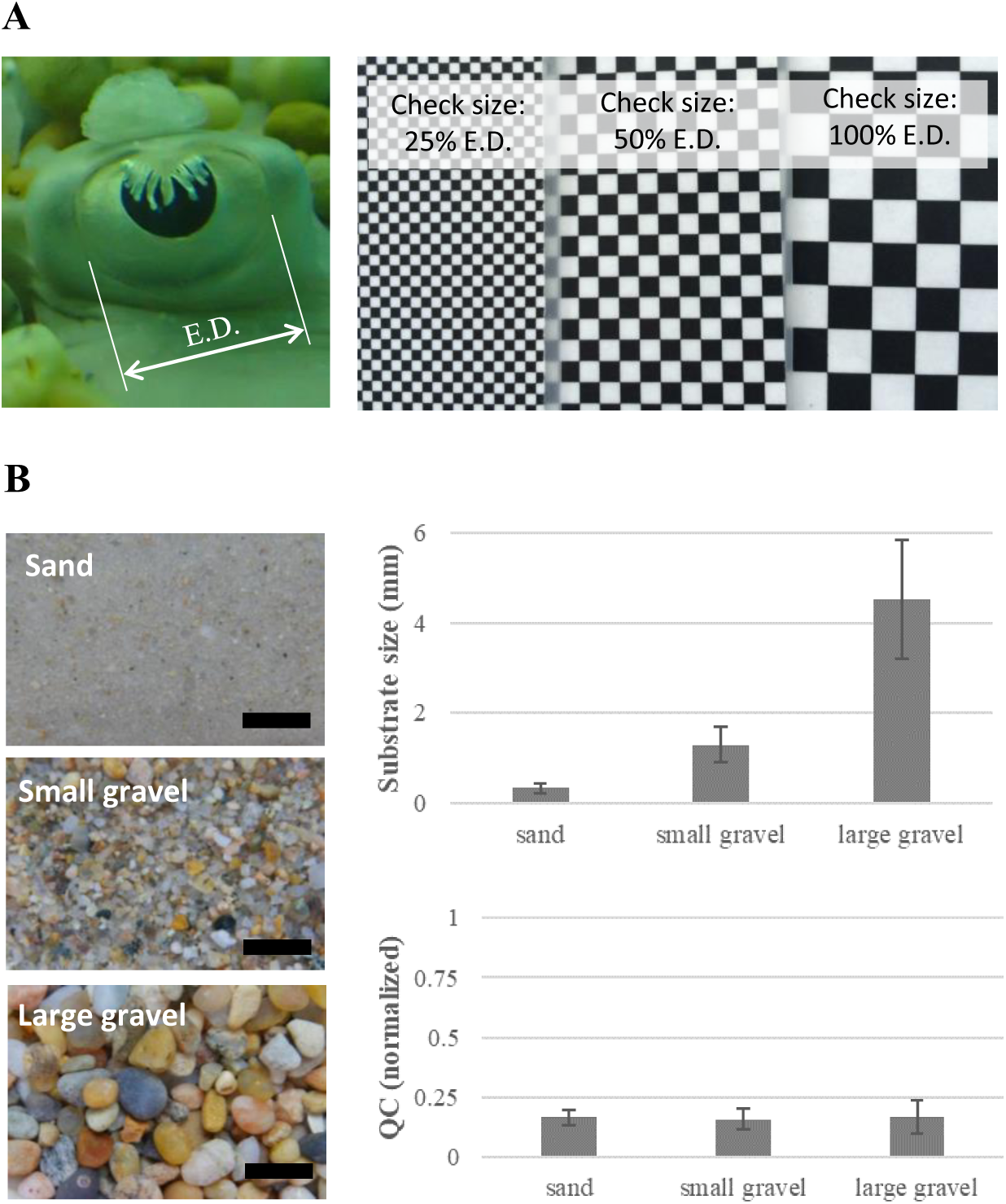
(A) Checkerboard substrates for Experiment 1 were created relative to skate eye dimensions. The largest checkerboard used a check size/length equivalent to the anterior-posterior dimension of the skate’s eye. The medium checkerboard’s size was 50% of the eye dimension; the small checkerboard 25% of the eye dimension. (B) Images of the natural substrates used in Experiment 2. Scale bars: 1 cm. Top right graph; substrate size (diameter of substrate objects, in mm). Bottom right panel; QC (quantum catch, a measure of the relative intensity at which an animal visually perceives a given object) of the three natural substrates used in Experiment 2 (Note, there was no statistical difference between substrates; see text for details).

Checkerboard substrates were placed on the bottom of the aquarium. We also placed checkerboards on the inside of the tank walls (checkerboard walls were 10 cm high and placed along the back and side walls, leaving the front wall open for filming). There was no particular order in which the skates were placed on their respective checkerboards. For each experiment, the lowest light intensity was set up first and filters were progressively removed to proceed to the next brighter light intensity (this proved more feasible than sliding filters into the light boxes to reduce light intensity). For each light setting, skates were given 30 minutes acclimation time, to ensure that the skate was calm, and resting on the aquarium floor, ready for the experiment. During this time, they were video recorded for 10 seconds every 5 minutes, so that the pupil dilation process could be documented. After 10-15 minutes, there was usually no change in pupil dilation. After a total of 30 minutes acclimation, the skates were filmed for 2 minutes. We then removed the neutral density filters to proceed to the next brighter light setting and allowed another 30 minutes adaptation, during which the pupil dilation was recorded as described before. After 30 minutes, the skate was video recorded for 2 minutes. This procedure was repeated for each light intensity.

The video camera was positioned horizontally, and perpendicular to the long axis of the skate (i.e., directly looking at the skate’s eye from the side), so that we were able to document the entire pupillary area during dilation and constriction (note, when the eye is seen from angles other than perpendicular to the skate’s long axis, the apparent total pupil area becomes smaller, so it was crucial to maintain this camera angle throughout the experiment). The camera angle was also set to be perpendicular to the aquarium glass to avoid error caused by diffraction. While filming, the filters on the outside of the tank were lifted slightly to accommodate the camera. Because of the tank dimensions, skates usually positioned themselves in such a way that the eye was perpendicular to the camera. This was determined by ensuring that both eyes (the eye that was being filmed, and the eye on the opposite side) were in a straight line, perpendicular to the camera. If the opposite eye was ahead or behind the eye being filmed, the skate was not aligned correctly. If skates had to be re-aligned, animals were generally agreeable to being gently moved and remained settled. Every skate was recorded when settled on its custom-made large, medium and small checkerboards at all three light intensities (0.68 l×, 10.9 l× and 43.75 l×).

*Experiment 2: Natural substrates experiment.* The experimental tank for this experiment measured 60 cm length, 60 cm width and 45 cm height, placed into the same dark room set-up described above. A larger tank was needed to allow skates to move about and bury themselves. Consequently, the overhead light source consisted of three custom-made light boxes (lined with LE Flexible Strip, SMD 2835 Daylight White LEDs) to provide even illumination across the entire tank (less than 5% variation; verified by Extech EasyView 30 light meter; at full illumination, the light intensity was measured to be approximately 500 l×). The tank was lined with the same filters as described for experiment 1. Neutral density filters (combination of 0.15ND and 0.6ND, Lee Filters) were used to obtain three light intensities that were previously verified to result in three distinct pupil dilation states: fully dilated, semi-constricted, near-constricted. These light intensities were 0.012 l×, 1.95 l×, 31.25 l×, respectively.

Three natural substrates were collected locally (Fig. 2B). These substrates are typical of the sandy/pebbly habitats in which skates are naturally found. ImageJ (NIH) was used to determine grain size (for each substrate, 100 particles were measured to determine average substrate size). (a) Sand, average grain size: 0.33 mm (0.11 s.d.); (b) Small gravel, average grain size: 1.30 mm (0.40 s.d.); (c) Large gravel, average grain size 4.5 mm (1.32 s.d.).

To ensure that skates responded to the spatial frequency of the natural substrates and not to variation on overall reflectance, we measured spectral reflectance of the three natural substrates using a spectrometer (QE65000, Ocean Optics) to ensure that overall reflectance was acceptably similar. A 200 µm fiber (400-UV-VIS, Ocean Optics) was held by a clip approximately 8 mm from the substrate at an angle of 30 degrees; the illumination consisted of the light source that was used for the behavioral trials (see above). A diffuse reflection standard (WS-1-SL, Ocean Optics), was used to standardize measurements. All measurements were taken in water. For each substrate, 70 measurements were taken. After the relative reflectance spectra were obtained, they were transformed into quantum units (divided by wavelength) and the number of photons (N) absorbed by a skate photoreceptor was calculated. This is given by

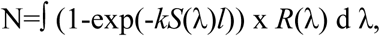

after Warrant (2004), where S(λ) is the spectral sensitivity of the visual pigment (500 nm; from Dowling and Ripps, 1970; Cornwall et al., 1989), R(λ) is the spectral composition of the light reflected from the substrate (both S(λ) and R(λ) integrated over 400 nm-650 nm), *l* is the length of the photoreceptor (67µm; from Ripps and Dowling, 1990) nd *k* is the quantum efficiency of transduction (0.037/µm; Warrant and Nilsson, 1998). The spectral sensitivity of the visual pigment was calculated using a template provided by A. Kelber, Lund, Sweden (based on Stavenga et al., 1993).

The three natural substrates yielded closely similar quantum catches; the following values are given as normalized values (on a scale of 0-1; 0 being the quantum catch for a pure black object, 1 being the quantum catch for a pure white object): large gravel average quantum catch: 0.167 (0.072 s.d.; ranges [min, max]: 0.046, 0.472); small gravel average quantum catch: 0.158 (0.046 s.d.; ranges [min, max]: 0.079, 0.328); sand average quantum catch: 0.164 (0.033 s.d.; ranges [min, max]: 0.072, 0.261) (Fig. 2B). A one-way ANOVA comparing the means of all three quantum catches (i.e., each of the substrates, as seen by a skate) revealed that there was no significant difference between the substrates [F(2,207)=0.54, p>0.5].

The bottom of the experimental tank was covered with substrate to a depth of approximately 5 cm. Skates were placed into the experimental tank set up at the lowest light intensity (0.012 l×). Compared to Experiment 1, skates needed a longer acclimation time to settle; thus, animals were given 50 minutes to acclimate. Next, skates were video-recorded for 2 minutes (same methods as used in experiment 1). Neutral density filters were then removed to obtain the next brightest light intensity (1.95 l×), followed by 50 minutes acclimation time and 2 minutes video recording. The same was repeated for the highest light intensity (31.25 l×). Once all skates were filmed at all light intensities on one substrate, we switched to the next substrate and repeated this procedure.

*(3) Analysis.* From the 2-minute video clips taken at each light intensity, we extracted 4-5 images at intervals of at least 10 seconds apart (since these were life animal experiments, we had to be flexible regarding the number of images we were able to obtain and the time between images). Using these images, we measured pupil dilation/constriction by determining pupil area (in mm^2^). This was done by tracing the exact outline of the pupil in ImageJ (NIH). To test for statistical significance, we used these pupil area measurements in a one-way ANOVA, with an alpha value of 0.05.

## Results

*Experiment 1: Checkerboard experiment.* For each light intensity, skates maintained almost the same pupil area, irrespective of check size (Fig. 3A). There was no statistical difference in pupil area on different checkerboards at each of the light intensities; one-way ANOVA - 0.68 l×: [F(2,117)=0.65, p>0.5]; 10.9 l×: [F(2,117)=0.34, p>0.7]; 43.75 l×: [F(2,117)=1.48, p>0.2]. Pupil area only changed in response to light intensity, with a significant difference between 0.68 l×, 10.9 l× and 43.75 l×. Since pupil area was the same on all check sizes irrespective of light intensity, all data for each light intensity were averaged, and a one-way ANOVA comparing the means on all three light intensities revealed that there was a significant difference in pupil area measured at different light intensities [F(2,357)=378.5, p<0.0001].

**Fig. 3.**
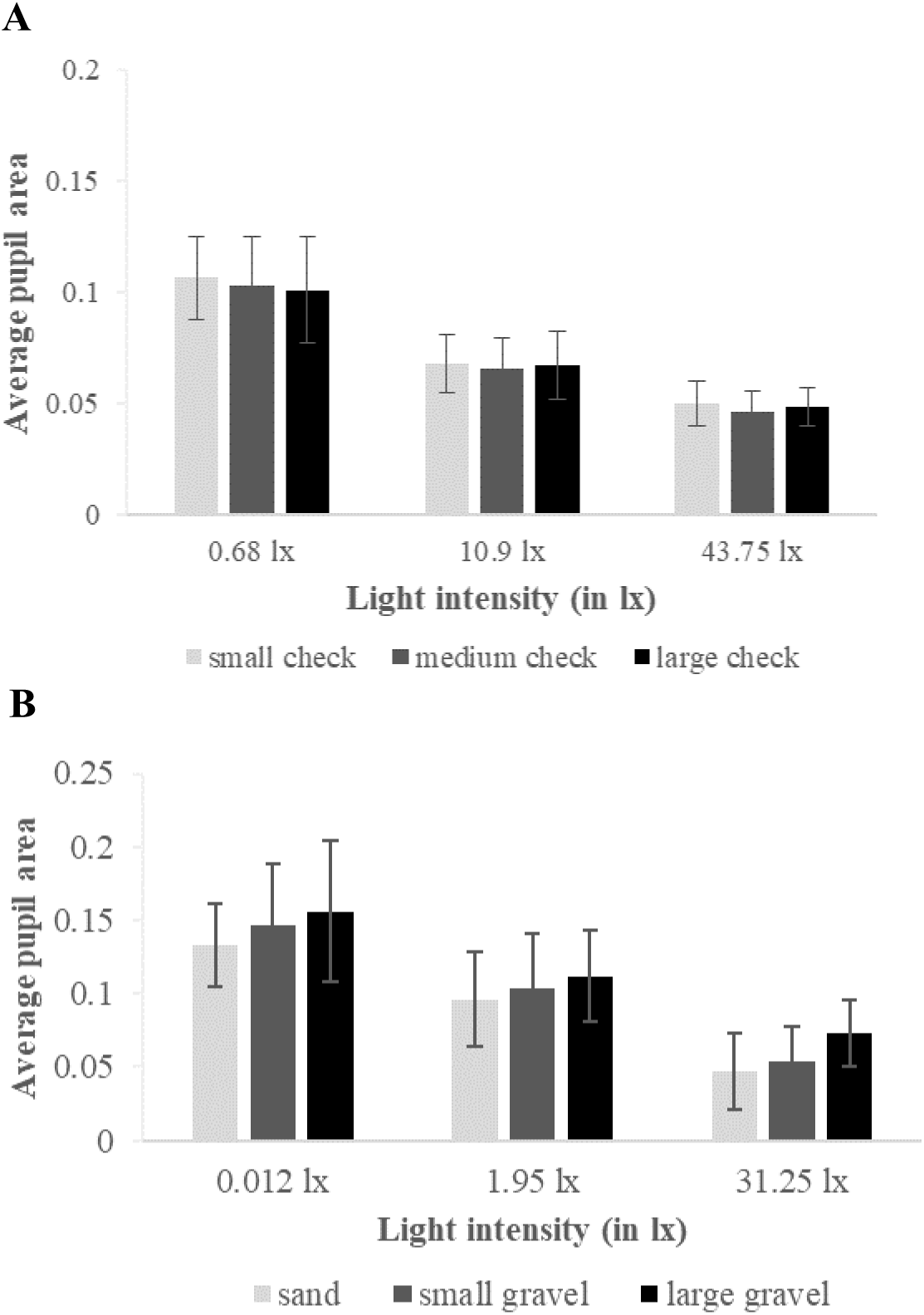
(A) Experiment 1: Pupil dilation for skates placed on artificial checkerboards. The skate pupil constricted with increasing light intensity but did not respond to changes in spatial frequency of the checkerboards (there was no significant difference between check sizes; there was a significant difference between light intensity; see text for details). (B) Experiment 2: Pupil dilation for skates allowed to bury in natural substrates, with only their eyes protruding. In addition to reacting to light intensity, the skate pupil constricted/dilated in response to the spatial frequency of the substrate: the smaller the substrate spatial frequency, the more constricted the pupil. This effect was statistically significant (see text for details). (Note, different dilation:constriction ratios in panels A and B are due to skate size differences).

*Experiment 2: Natural substrates experiment.* As in Experiment 1, pupil area changed in response to light intensity, with a significant difference between 0.012 l×, 1.95 l×, and 31.25 l× (one-way ANOVA; sand [F(2, 165)=127.7, p<0.0001; small gravel [F(2, 165)=100.4, p<0.0001; large gravel [F(2, 165)=80.3, p<0.0001]). In contrast to Experiment 1, skate pupil dilation also depended on substrate spatial frequency (Fig. 3B). Given the same light intensity, the pupils were more constricted on substrates with smaller spatial frequency, and more dilated on substrates with larger spatial frequency. This effect was statistically significant (one-way ANOVA; 0.012 l×: [F(2, 165)=4.98, p=0.0078]; 1.95 l×: [F(2, 165)=3.2, p=0.043]; 31.25 l×: [F(2,165)=17.56, p<0.0001).

## Discussion

While it has been suggested that pupils may aid in camouflaging the eye (Muntz, 1977; Douglas 2018; Douglas et al., 1998; 2002; 2005), this has never been tested experimentally. The two experiments reported here present evidence that the elaborate pupil shape of the skate *Leucoraja erinacea* may indeed serve a camouflage function. When skates were placed on artificial backgrounds of different spatial frequencies (checkerboards with different check sizes), their pupils only dilated/constricted in response to light intensity – not check size. When placed on natural substrates that allowed burying, so that only the eyes protruded, the skates’ pupils dilated/constricted in response to the spatial frequency of the substrate as well as light intensity. While these findings seemingly contradict each other, when considered in the context of how camouflage works, the possible rationality behind this quickly emerges. Object recognition is subject to a multi-step process that begins with the detection of the object by the observer’s visual system. This depends on the visual system of the observer, the physical characteristics of the object, and the background, against which the object is seen. Background characteristics are obviously paramount in the context of camouflaged objects (Thayer, 1909; Longley, 1916; Longley, 1917; Cott, 1940; Endler, 1981, 1984; Crook, 1997a,b; Barry and Hawryshyn, 1999; Marshall, 2000; Vorobyev et al., 2001; Marshall et al., 2003). Going back to experiment 1, in which the skates were placed on artificial backgrounds that did not permit burying, the background against which an observer would see the skates’ eyes was not the checkerboard substrate but the skate’s head and body (because the skate was not buried into the substrate). Consequently, unless the skate’s head and body colors or patterns change in response to the spatial frequencies of the checkerboard substrates, there may be no reason for why the skate’s pupil would be expected to react to the different background spatial frequencies. Skates do not have the fast color changing abilities we see in some teleost fish and cephalopods (Parker and Porter, 1934; Waring, 1963; Hanlon and Messenger, 2018), and, while elasmobranchs are capable of subtle color tone and pattern changes (Parker and Porter, 1934; Parker, 1936; Visconti et al., 1999), these changes generally occur over the course of many hours. The short duration of our experiments was likely not long enough for any color or pattern changes to occur. Notably, we did not observe any color or pattern changes in our experiments.

In Experiment 2, burying behavior was permitted, and all skates buried themselves. In this situation, the eyes were seen against the natural substrate, rather than the skate’s body; thus, the fact that the skate’s pupil changed with changes in background spatial frequency is entirely rational.

In order to understand the possible rationale behind the skate pupillary movements we report here, it is necessary to briefly mention some fundamental camouflage principles. A range of colors and patterns have evolved in many animals for the sole purpose of reducing the risk of detection, with the ultimate goal of evading predation (e.g., Endler, 1978; Stevens and Cuthill, 2006; Stevens and Merilaita, 2009; Troscianko et al., 2009; Stevens and Merilaita, 2011). To avoid detection, several cryptic mechanisms are deployed by animals: briefly, they include background matching, self-shadow concealment, obliterative shading, disruptive coloration, flicker-fusion camouflage, and distractive markings (Stevens and Merilaita, 2009). While avoiding detection would clearly be the preferred option, this is often hard to achieve in nature. Animals move between habitats and are therefore frequently faced with changing background characteristics, yet most animals have only one, or a limited number of body colors and patterns available to them. Additionally, some structures - e.g., the eyes - are impossible to conceal to a level that would completely eliminate detection: the pupil, by definition, is a dark hole for light to enter so that vision can ensue. As a result, many animals have solved this problem by having physical characteristics that reduce the likelihood of recognition, for example, by creating false contours or distractive markings, or by disguising the body’s true outline by disruptive coloration (Cuthill and Székely, 2009). These forms of camouflage are aimed at fooling a specific observers’ perceptual processes, for example their visual system’s color or polarization vision ability, or their edge detection mechanisms (Stevens and Merilaita, 2009; Troscianko et al., 2009; Stevens and Merilaita, 2011). It is likely that the camouflage mechanisms underlying the elaborate pupil shape of the skate falls under this category; specifically, the irregular pupil outline creates an edge pattern that, compared to a conventional circular pupil pattern, would be much harder to detect. Since skates like to bury into the substrate to practice their various predatory life styles (active foraging, sit-and-wait tactics), the skate eye is usually the only part of the animal that is visible. The iris structures that form their elaborate pupil shapes are lined with silvery reflectors, similar to the ones we see in cephalopod eyes (Denton and Land, 1971; Mäthger et al., 2009), providing the animal with the ability to match the surrounding natural background in color and brightness. However, as discussed in illustrative detail by Cott (1940), animal eyes are intrinsically difficult to conceal because of their highly regular structures that do not blend well into the usually irregular spatial features of natural backgrounds. Elaborate pupil shapes, such as those of skates, may therefore aid camouflage by introducing irregular shapes and lines that break up the otherwise conspicuous outline of the pupil, so that it may reduce the likelihood of being recognized as part of an eye that belongs to a potential predator (or prey). Additionally, to help achieve the required level of camouflage on a range of natural substrates, it would seem advantageous to adjust pupil dilation and constriction relative to background spatial frequency, which is what we found in this study. However; in skates, these dilation and constriction changes were small, and did not exactly match the spatial frequencies of the surrounding backgrounds. More work is clearly needed to fully evaluate the contribution of pupillary movements in camouflage on backgrounds of different spatial frequencies. For example, the size ranges of the natural and artificial substrates did not completely overlap. The artificial checkerboard substrates concentrated on higher - whereas the natural substrates concentrated on lower spatial frequencies. It would be valuable to study the pupillary response at the lower spatial frequency range, which is more relevant to these animals’ life style.

In conclusion, while a crucial purpose of the pupil is likely to regulate the amount of light entering the eye, a slight departure from the pupil’s optimal dilation/constriction state to a given light intensity can be tolerated by the retina. By compromising retinal illumination, the animal may gain a critical camouflage advantage.

## Acknowledgements

SY and CO were undergraduate student interns. The authors would like to thank Dan-Eric Nilsson for stimulating discussions, and the staff of the Marine Resources Center at the MBL for assistance with skates. This work was funded by an award from the Marine Biological Laboratory (Hermann Foundation Award, Joan Ruderman Fund Award, Grass Foundation Fund Award, Neal Cornell Career Development Award), as well as a University of Chicago Metcalf fellowship to CO.

## Author contributions

LMM conceived the study, assisted with data collection, analyzed data and wrote the manuscript. SY and CO (both undergraduate student interns) collected data and assisted with data analysis and manuscript preparation (both authors contributed equally).

